# The MADS-box transcription factor PHERES1 controls imprinting in the endosperm by binding to domesticated transposons

**DOI:** 10.1101/616698

**Authors:** Rita A. Batista, Jordi Moreno-Romero, Yichun Qiu, Joram van Boven, Juan Santos-González, Duarte D. Figueiredo, Claudia Köhler

## Abstract

MADS-box transcription factors are ubiquitous in eukaryotic organisms and play major roles during plant development. Nevertheless, their function in seed development remains largely unknown. Here we show that the imprinted *Arabidopsis thaliana* MADS-box TF PHERES1 (PHE1) is a master regulator of paternally expressed imprinted genes, as well as of non-imprinted key regulators of endosperm development. PHE1 binding sites show distinct epigenetic modifications on maternal and paternal alleles, correlating with parental-specific transcriptional activity. Importantly, we show that the CArG-box-like DNA-binding motifs bound by PHE1 have been distributed by RC/Helitron transposable elements. Our data provide an example of molecular domestication of these elements, which by distributing PHE1 binding sites throughout the genome, have facilitated the recruitment of crucial endosperm regulators into a single transcriptional network.

## Introduction

MADS-box transcription factors (TFs) are present in most eukaryotes, and classified into two groups: type I or SRF (Serum Response Factor)-like, and type II or MEF2 (Myocyte Enhancing Factor2)-like (Gramzow and Theissen, 2010). In flowering plants, type I MADS-box TFs are associated with reproductive development and many are active in the endosperm, a nutritive seed tissue that supports embryo growth (Bemer et al., 2010). Deregulation of type I MADS-box TF genes has been frequently linked to failure of endosperm development and seed inviability, both in the Brassicaceae (Erilova et al., 2009; Lu et al., 2012; Rebernig et al., 2015; Tiwari et al., 2010), and in crop species like tomato and rice (Ishikawa et al., 2011; Roth et al., 2019). In *Arabidopsis thaliana* and other species, the activity of type I MADS-box TFs has been shown to be associated with the timing of endosperm cellularisation – an important developmental transition, which is fundamental for seed viability (Erilova et al., 2009; Kang et al., 2008; Lu et al., 2012; Tiwari et al., 2010). Nevertheless, a mechanistic explanation for these observations, clarifying the role of MADS-box TFs in endosperm development, remains to be established.

In this work we characterise the function of the type I MADS-box TF PHE1. *PHE1* is active in the endosperm and is a paternally expressed imprinted gene (Köhler et al., 2005). Imprinting is defined as an epigenetic phenomenon causing a gene to be preferentially expressed from the maternal or the paternal allele. In plants, imprinting is mainly manifested in the endosperm, and similarly to type I MADS-box TFs, imprinted genes have been previously implicated in endosperm development (Erilova et al., 2009; Figueiredo et al., 2015; Jullien and Berger, 2010; Pignatta et al., 2018; Wolff et al., 2015). Imprinting relies on the asymmetric distribution of epigenetic marks in maternal and paternal alleles, which is thought to distinctly impact their transcription (Gehring, 2013; Rodrigues and Zilberman, 2015). However, transcriptional regulators of imprinted genes have not been identified until now. Thus, the interaction between epigenetic modifications of imprinted genes and their transcriptional regulation remains unknown. Our results identify PHE1 as a key transcriptional regulator of imprinted genes, as well as of other genes important for endosperm development. Furthermore, we explore the cross-talk between epigenetic regulation and PHE1 transcriptional activity at imprinted gene loci, and show that differential parental epigenetic modifications of PHE1 binding sites correlates with the transcriptional status of the parental alleles. Finally, we reveal that RC/Helitron transposable elements (TEs) have served as distributors of PHE1 DNA-binding sites, providing an example of molecular domestication of TEs.

## Results and Discussion

To identify the genes regulated by PHE1, we performed a ChIP-seq experiment using siliques from a PHE1::PHE1-GFP line, which has been previously shown to expressed PHE1-GFP exclusively in the endosperm (Weinhofer et al., 2010). We identified a total of 1942 PHE1 target genes, which were enriched for Gene Ontology (GO)-terms associated with development, metabolic processes, and transcriptional regulation, showing that many PHE1 targets are themselves transcriptional regulators (Fig. 1 – Fig. Supplement 1a). Among PHE1 targets are several known regulators of endosperm development like *AGL62, YUC10, IKU2, MINI3*, and *ZHOUPI*, revealing a central role for PHE1 in regulating this developmental process (Fig. 1 – Fig. Supplement 2). Importantly, type I MADS-box family genes were over-represented among PHE1 targets (Fig. 1 – Fig. Supplement 1a), pointing to a high degree of cross-regulation among members of this family.

Our data revealed that PHE1 uses two distinct DNA-binding motifs: motif A was present in about 53% of PHE1 binding sites, while motif B could be found in 43% of those (Fig. 1a). PHE1 motifs closely resemble type II CArG-boxes, the signature motif of MADS-box TFs (Folter and Angenent, 2006) (Fig. 1a, Fig. 1 – Fig. Supplement 3), suggesting that DNA-binding properties between type I and type II members are conserved.

**Figure 1.**
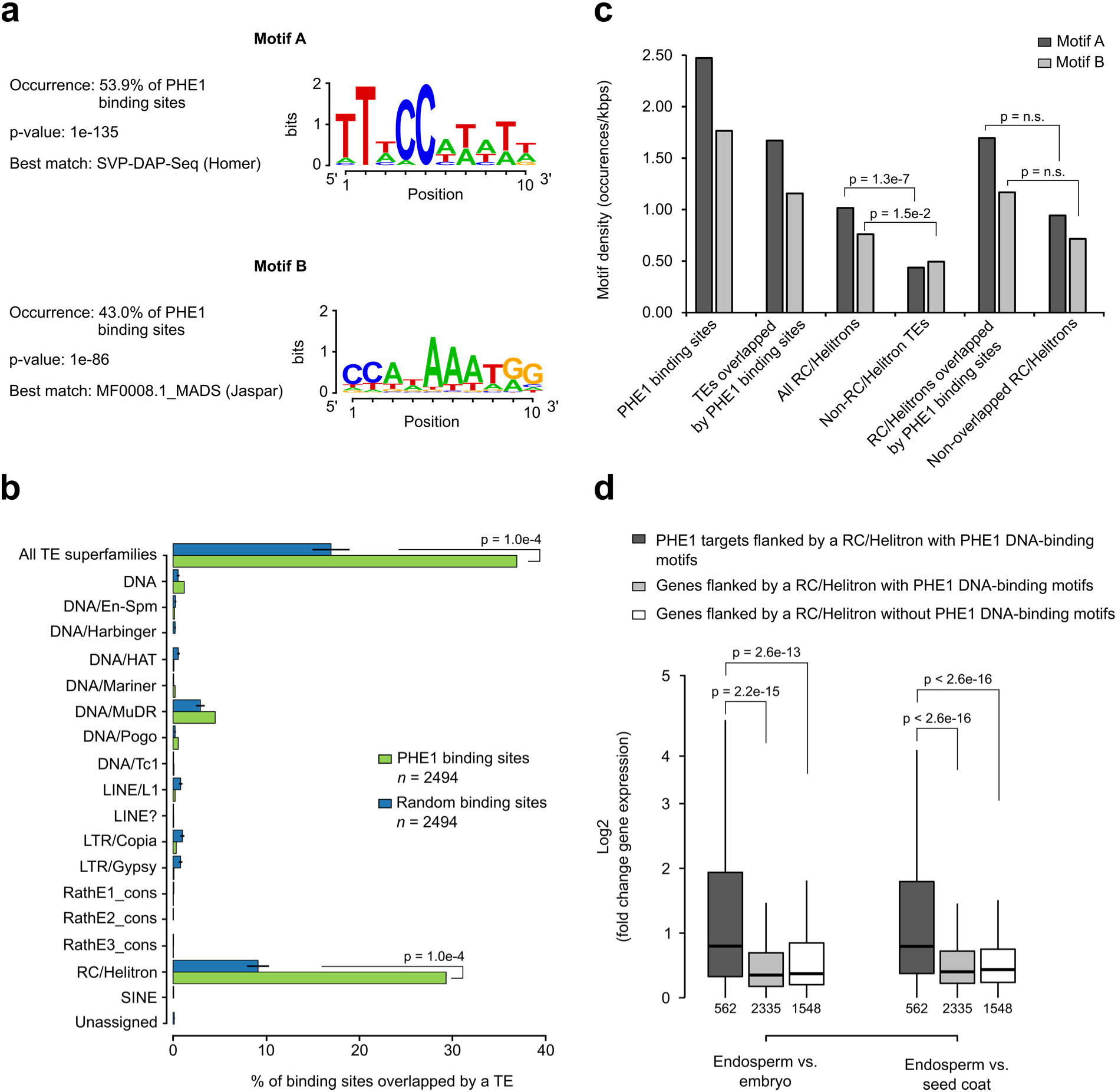
RC/Helitrons carry PHE1 DNA-binding motifs. **a,** CArG-box like DNA-binding motifs identified from PHE1 ChIP-seq data. **b,** Fraction of PHE1 binding sites (green) overlapping transposable elements (TEs.) Overlap is expressed as the percentage of total binding sites where spatial intersection with features on the y-axis is observed. A set of random binding sites is used as control (blue). This control set was obtained by randomly shuffling the PHE1 binding sites in *A. thaliana* gene promoters (see Methods). P-values were determined using Monte Carlo permutation tests (see Methods). Bars represent ± sd, (n = 2494, PHE1 binding sites, Random binding sites). **c,** Density of PHE1 DNA-binding motifs in different genomic regions of interest. P-values were determined using χ2 tests. **d,** Fold change gene expression of genes flanked by RC/Helitron TEs. Fold change was determined between endosperm and embryo, or between endosperm and seed coat. Genes were divided in three categories depending on their PHE1 target status, and the presence of RC/Helitrons with and without PHE1 DNA-binding motifs. Gene expression data was retrieved from Belmonte et al., (2013). Pre-globular seed stage was used in this analysis. P-values were determined using two-tailed Mann-Whitney tests (*n =* number represented below boxplots).

Strikingly, PHE1 binding sites significantly overlapped with TEs, preferentially with those of the RC/Helitron superfamily (29% (723/2494) in PHE1 binding sites, versus 9% (224/2494) in random binding sites) (Fig. 1b), and in particular with some RC/Helitron subfamilies (Fig. 1 – Fig. Supplement 4). We thus addressed the question whether RC/Helitrons contain sequence properties that promote PHE1 binding. Indeed, a screen of genomic regions for the presence of PHE1 DNA-binding motifs, revealed significantly higher motif densities within all RC/Helitrons, compared to other TE superfamilies (Fig. 1c). We found that most RC/Helitron sequences associated with both PHE1 DNA-binding motifs share sequence homology, and the majority of them could be grouped into one large cluster, based on sequence identity (Fig. 1 – Figure Supplement 5). While the presence of additional smaller clusters points to a few instances of independent gains of PHE1 DNA-binding motifs through *de novo* mutation or sequence capture, the grouping of most sequences within one cluster suggests that a single ancestral RC/Helitron likely acquired a perfect or nearly perfect motif sequence. Moreover, we could detect the presence of perfect or nearly perfect PHE1 DNA-binding motifs within the consensus sequences of several RC/Helitron families (Fig. 1 – Figure Supplement 6), suggesting that radiation of RC/Helitrons occurred after the acquisition of the binding motif. Together, these data suggest that an ancestral RC/Helitron likely played a role in acquiring PHE1 DNA-binding motifs, which were subsequently amplified in the genome through transposition.

Interestingly, even though motif densities were higher in RC/Helitrons overlapped by PHE1 binding sites than in non-overlapped RC/Helitrons, this difference was not significant (Fig. 1c). As the enrichment of PHE1 DNA-binding motifs is a specific feature of RC/Helitrons, the domestication of these TEs as *cis*-regulatory regions might facilitate TF binding and modulation of gene expression. In line with this, we detected that genes flanked by RC/Helitrons carrying bound PHE1 DNA-binding motifs were more highly expressed in the endosperm than in other seed tissues (Fig. 1d). This shows that for a subset of genes, these TEs can be effectively used as sites for PHE1 binding, thus triggering endosperm-specific expression.

Among the PHE1 target genes we detected a significant enrichment of imprinted genes, with 12% of all maternally expressed genes (MEGs) and 31% of all paternally expressed genes (PEGs) being targeted (Fig. 2a). Imprinted gene expression relies on parental-specific epigenetic modifications, which are asymmetrically established during male and female gametogenesis, and inherited in the endosperm (Gehring, 2013; Rodrigues and Zilberman, 2015). Demethylation of repeat sequences and TEs – occurring in the central cell, but not in sperm – is a major driver for imprinted gene expression (Gehring, 2013; Rodrigues and Zilberman, 2015). In MEGs, DNA hypomethylation of maternal alleles leads to their expression, while DNA methylation represses the paternal allele (Gehring, 2013; Rodrigues and Zilberman, 2015). In PEGs, the hypomethylated maternal allele undergoes trimethylation of lysine 27 of histone H3 (H3K27me3), a repressive histone modification established by the Fertilization Independent Seed-Polycomb Repressive Complex2 (FIS-PRC2). This renders the maternal alleles inactive, while the paternal allele is expressed (Gehring, 2013; Moreno-Romero et al., 2016; Rodrigues and Zilberman, 2015).

**Figure 2.**
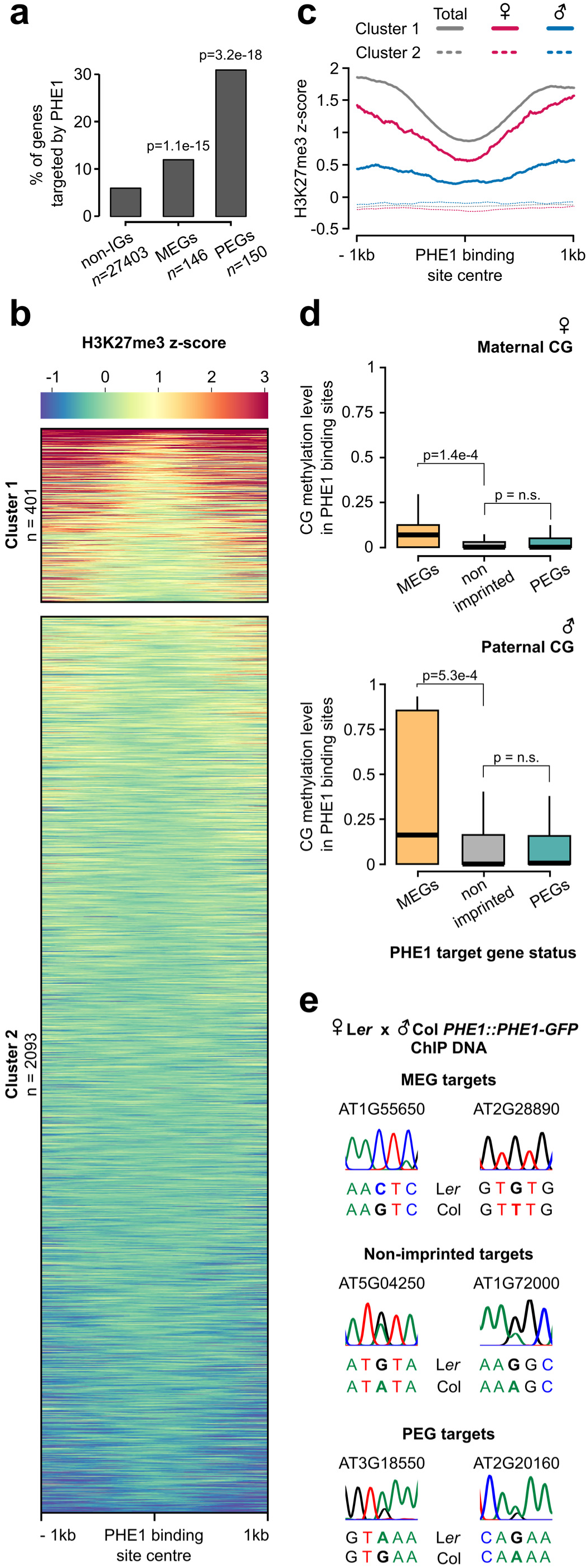
Parental asymmetry of epigenetic marks in imprinted gene promoters conditions PHE1 binding. **a,** Fraction of non-imprinted and published imprinted genes targeted by PHE1. P-values were determined using the hypergeometric test. The list of published imprinted genes used for this analysis is detailed in **Fig 1. - source data 1**. **b,** Heatmap of endosperm H3K27me3 distribution along PHE1 binding sites. Each horizontal line represents one binding site. Clusters were defined based on the pattern of H3K27me3 distribution (see Methods) **c,** Metagene plot of average maternal (♀, pink), paternal (♂, blue) and total (grey) endosperm H3K27me3 along PHE1 binding sites. **d,** CG methylation levels in maternal (♀, upper panel) and paternal (♂, lower panel) alleles of PHE1 binding sites associated with MEGs (yellow), PEGs (green) and non-imprinted (grey) PHE1 targets. P-values were determined using two-tailed Mann-Whitney tests. **e,** Sanger sequencing of imprinted and non-imprinted gene promoters bound by PHE1. SNPs for maternal (L*er*) and paternal (Col) alleles are shown, (*n =* 1 biological replicate). Maternal:total read ratios for imprinted genes are as follows: AT1G55650 – 1.0; AT2G28890 – 0.97; AT3G18550 – 0.44; AT2G20160 – 0.32.

Given the significant overrepresentation of imprinted genes among PHE1 targets, we assessed how the epigenetic landscape at those loci correlates with DNA-binding by PHE1. We surveyed levels of endosperm H3K27me3 within PHE1 binding sites and identified two distinct clusters (Fig. 2b). Cluster 1 was characterized by an accumulation of H3K27me3 in regions flanking the centre of the binding site, while the centre itself was devoid of this mark. Cluster 2 contained binding sites largely devoid of H3K27me3. The distribution of H3K27me3 in cluster 1 was mostly attributed to the deposition of H3K27me3 on the maternal alleles, while the paternal alleles were devoid of this mark (Fig. 2c) – a pattern usually associated with PEGs (Moreno-Romero et al., 2016). Consistently, genes associated with cluster 1 binding sites had a more paternally-biased expression in the endosperm when compared to genes associated with cluster 2 (Fig. 2 – Fig. Supplement 1a). This is reflected by more PEGs and putative PEGs being associated with cluster 1 (Fig. 2 – Fig. Supplement 1b). We also identified parental-specific differences in DNA methylation, specifically in the CG context: PHE1 binding sites associated with MEGs had significantly higher methylation levels in paternal alleles than in maternal alleles (Fig. 2d).

We hypothesized that the differential parental deposition of epigenetic marks in parental alleles of PHE1 binding sites can result in differential accessibility of each allele. This might impact the binding of PHE1, and therefore also impact transcription in a parent-of-origin-specific manner. To test this, we performed ChIP using a L*er* maternal plant and a Col *PHE1::PHE1-GFP* pollen donor, taking advantage of SNPs between these two accessions to discern parental preferences of PHE1 binding. Using Sanger sequencing, we determined the parental origin of enriched ChIP-DNA in MEG, non-imprinted, and PEG targets (Fig. 2e, Fig. 2 – Fig. Supplement 2). While binding of PHE1 was biallelic in non-imprinted targets (Fig 2e), only maternal binding was detected in the tested MEG targets, supporting the idea that CG hypermethylation of paternal alleles prevents their binding by PHE1 (Fig. 2e, Fig. 2 – Fig. Supplement 3a). Interestingly, we observed biallelic binding in PEG targets (Fig. 2e). Even though the maternal PHE1 binding sites in PEGs were flanked by H3K27me3 (Fig. 2b-c), correlating with transcriptional repression of maternal alleles, the absence of this mark within the binding site centres seems to be permissive for maternal PHE1 binding. We speculate that the accessibility of this site might be important to mediate recruitment of H3K27me3 in the central cell and/or for maintenance of this mark during endosperm development (Fig. 2 – Fig. Supplement 3a).

Previous studies have shown that PEGs are often flanked by RC/Helitrons (Pignatta et al., 2014; Wolff et al., 2011), a phenomenon suggested to lead to the parental asymmetry of epigenetic marks in these genes (Moreno-Romero et al., 2016; Pignatta et al., 2014). Consistent with our finding that PHE1 binding sites overlapped with RC/Helitrons (Fig. 1b), we found that PHE1 DNA-binding motifs were contained within these TEs significantly more frequently in PEGs than in non-imprinted genes (Fig. 2 – Fig. Supplement 3b). Furthermore, we detected the presence of homologous RC/Helitrons containing PHE1 binding motifs in the promoter regions of several PHE1-targeted PEG orthologs (Fig. 3 a-e, Fig. 3 – Figure Supplement 1), indicating ancestral insertion events. The presence of these RC/Helitrons correlated with paternally-biased expression of the associated orthologs, providing further support to the hypothesis these TEs contribute to the gain of imprinting, especially of PEGs. Thus, besides facilitating the asymmetry of epigenetic marks, these TEs can contribute to the generation of novel gene promoters that ensure the timely endosperm expression of PEGs, under the control of PHE1, and possibly other type I MADS-box TFs (Fig. 2 – Fig. Supplement 3a).

**Figure 3.**
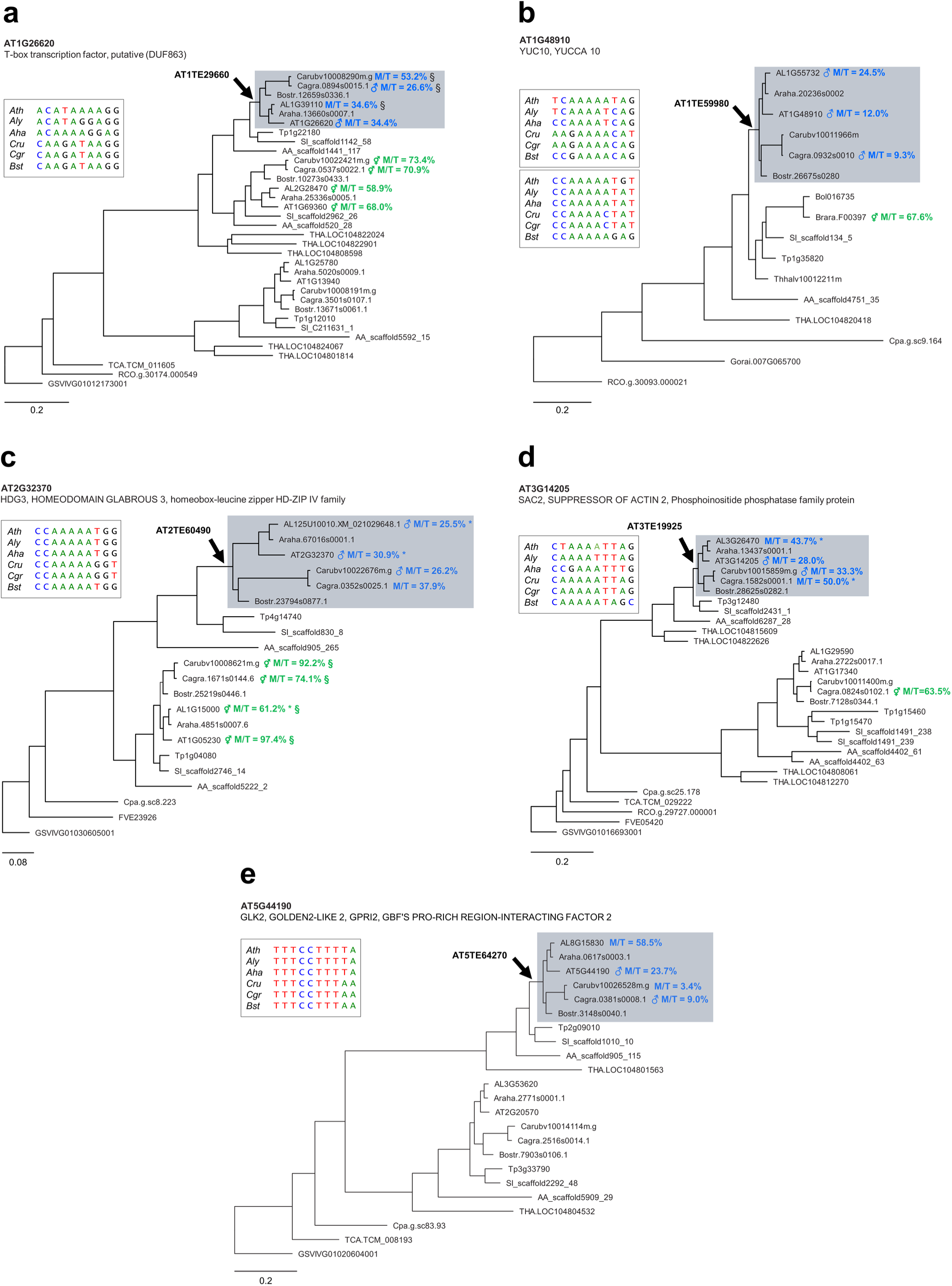
Ancestral RC/Helitron insertions are associated with gain of imprinting in the Brassicaceae. **a-e**, Phylogenetic analyses of PHE1-targeted PEGs and their homologs. Each panel represents a distinct target gene, and its corresponding homologs in different species. The genes in grey background have homologous RC/Helitron sequences in their promoter region. The arrow indicates the putative insertion of an ancestral RC/Helitron. The identity of the RC/Helitron identified in *A. thaliana* is indicated. These *A. thaliana* RC/Helitrons contain a PHE1-DNA binding motif and are associated with a PHE1 binding site. The inset boxes represent the alignment between the *A. thaliana* PHE1 DNA-binding motif and similar DNA motifs contained in RC/Helitrons present at the promoter regions of orthologous genes. When available, the imprinting status of a given gene is indicated by the presence of ♂ (PEG), or ⚥ (non-imprinted), and reflects the original imprinting analyses done in the source publications (see Methods). The maternal:total read ratio (M/T) for each gene is also indicated. §: potential contamination from maternal tissue. *: accession-biased expression. Scale bar represents substitution per site for the ML tree. The tree is unrooted. Gene identifier nomenclatures: AT, *Arabidopsis thaliana*; AL, *Arabidopsis lyrata*; Araha, *Arabidopsis halleri*; Bostr, *Boechera stricta*; Carubv, *Capsella rubella*; Cagra, *Capsella grandiflora*; Tp, *Schrenkiella parvula*; SI, *Sisymbrium irio*; Bol, *Brassica oleracea*; Brapa, *Brassica rapa*; Thhalv, *Eutrema salsugineum*; AA, *Aethionema arabicum*; THA, *Tarenaya hassleriana*; Cpa, *Carica papaya*; TCA, *Theobroma cacao*; Gorai, *Gossypium raimondii*; RCO, *Ricinus communis*; FVE, *Fragaria vesca*; GSVIVG, *Vitis vinifera*.

Among the PEGs targeted by PHE1 were *ADM*, *SUVH7*, *PEG2,* and *NRPD1a* (Fig. 1 – Fig. Supplement 2). Mutants in all four PEGs suppress the abortion of triploid (3x) seeds generated by paternal excess interploidy crosses (Martinez et al., 2018; Satyaki and Gehring, 2019; Wolff et al., 2015). Furthermore, we found that close to 50% of highly upregulated genes in 3x seeds are targeted by PHE1 (Fig. 4a), suggesting this TF might play a central role in mediating the strong gene deregulation observed in these seeds. If this is true, removal of PHE1 in 3x seeds is expected to suppress 3x seed inviability. To test this hypothesis, we generated a *phe1* CRISPR/Cas9 mutant in the *phe2* background, since both genes are likely redundant (Villar et al., 2009) (Fig. 4 – Fig. Supplement 1a). We introduced *phe1 phe2* into the *omission of second division 1 (osd1)* mutant background, which produces diploid gametes at high frequency (d’Erfurth et al., 2009). Wild-type (wt) and *phe2* maternal plants pollinated with *osd1* pollen form 3x seeds that abort at high frequency (Kradolfer et al., 2013) (Fig. 4b, Fig. 4 – Fig. Supplement 1b). In contrast, *phe1 phe2 osd1* pollen strongly suppressed 3x seed inviability, reflected by the increased germination of 3x *phe1 phe2* seeds (Fig. 4c, Fig. 4 – Fig. Supplement 1c). This phenotype could be reverted by introducing the *PHE1::PHE1-GFP* transgene paternally (Fig. 4b-c, Fig. 4 – Fig. Supplement 1b-c). Notably, 3x seed rescue was mostly mediated by *phe1*, as the presence of a wt *PHE2* allele in 3x seeds (wt x *phe1 phe2 osd1*) led to comparable rescue levels than when having no wt *PHE2* allele present (*phe2* x *phe1 phe2 osd1*) (Fig. 4b-c, Fig. 4 – Fig. Supplement 1b-c). Importantly, 3x seed rescue was accompanied by reestablishment of endosperm cellularisation (Fig. 4 – Fig. Supplement 1d-e), and reduced expression of PHE1 target genes (Fig. 4 – Fig. Supplement 1f).

**Figure 4.**
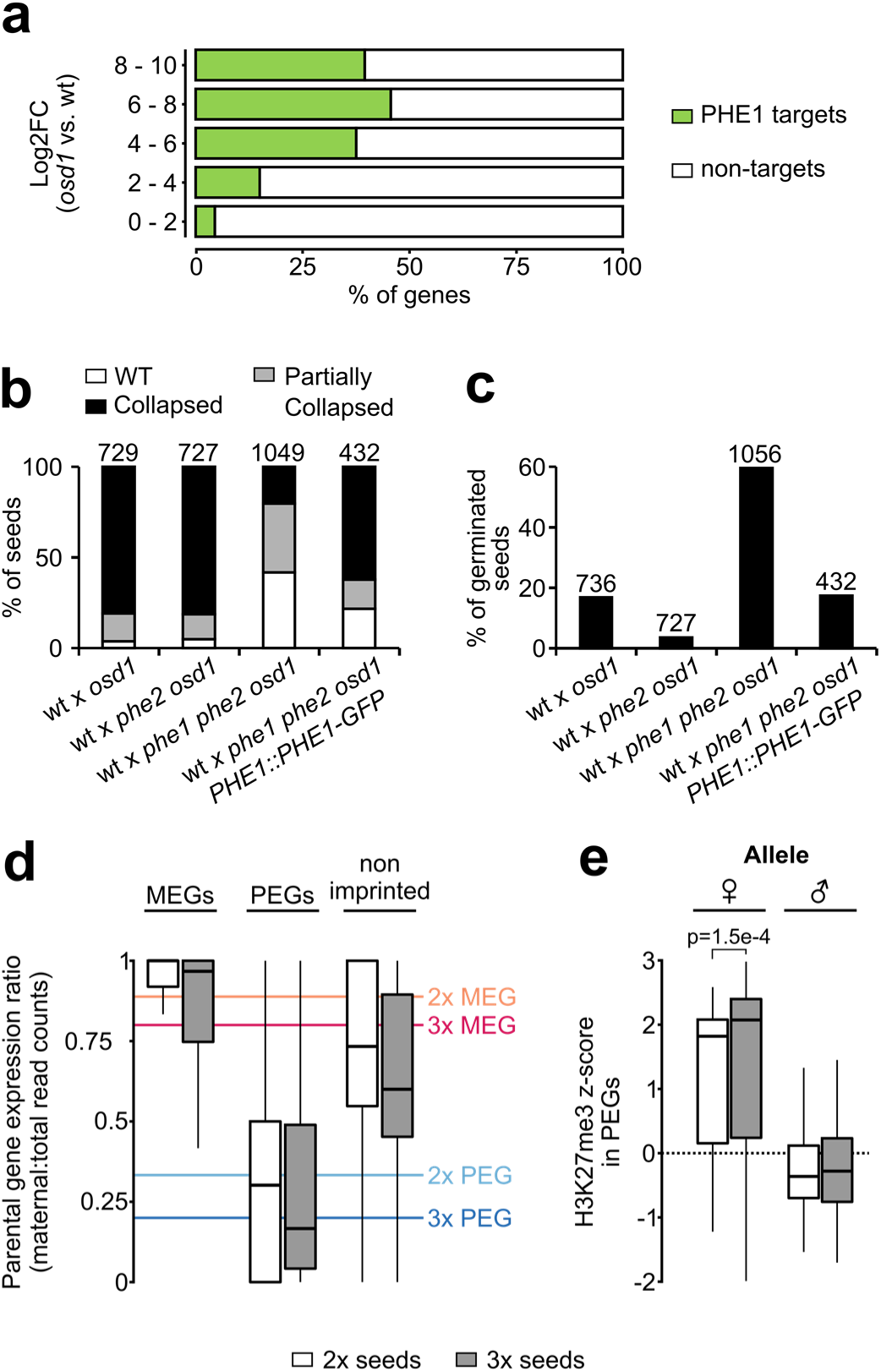
PHE1 establishes 3x seed inviability of paternal excess crosses. **a,** Target status of upregulated genes in paternal excess crosses. Highly upregulated genes in 3x seeds are more often targeted by PHE1 (p<2.2e-16, χ^2^ test). **b-c,** Seed inviability phenotype (b) of paternal excess crosses in wild-type (wt), *phe1 phe2*, and *phe1* complementation lines, with respective seed germination rates (c). The maternal parent is always indicated first. Remaining control crosses are shown in Fig. 4 – Fig. Supplement 1b-c. *n =* numbers on top of bars (seeds). **d,** Parental expression ratio of imprinted genes in the endosperm of 2x (white) and 3x seeds (grey). Solid lines indicate the ratio thresholds for definition of MEGs and PEGs, in 2x and 3x seeds. **e,** Accumulation of H3K27me3 across maternal (♀) and paternal (♂) gene bodies of PEGs in the endosperm of 2x and 3x seeds (white and grey, respectively). H3K27me3 accumulation in MEGs and non-imprinted genes is shown in Fig. 4 – Fig. Supplement 2. P-value was determined using a two-tailed Mann-Whitney test

Loss of FIS-PRC2 function causes a similar phenotype to that of paternal excess 3x seeds, correlating with largely overlapping sets of deregulated genes (Erilova et al., 2009; Tiwari et al., 2010). Since FIS-RC2 is a major regulator of PEGs in *Arabidopsis* endosperm (Moreno-Romero et al., 2016), we addressed the question whether imprinting is disrupted in 3x seeds. To assess this, we analysed the parental expression ratio of imprinted genes in the endosperm of 2x and 3x seeds. Surprisingly, imprinting was not disrupted in 3x seeds (Fig. 4d), consistent with the presence of similar H3K27me3levels on the maternal alleles of PEGs in the endosperm of 2x and 3x seeds (Fig. 4e, Fig. 4 – Fig. Supplement 2). Collectively, these data show that the major upregulation of imprinted gene expression in 3x seeds is due to increased transcription of the active allele, likely mediated by PHE1 and other MADS-box TFs, with maintenance of the imprinting status.

In summary, this work reveals that the MADS-box TF PHE1 is a major regulator of imprinted genes in the *Arabidopsis* endosperm, and that this TF establishes a reproductive barrier in response to interploidy hybridizations. We furthermore show that deregulated PEGs in 3x seeds remain imprinted, but that the active allele becomes strongly overexpressed, correlating with increased PHE1 activity in these seeds (Erilova et al., 2009). Importantly, we reveal a novel role for RC/Helitrons in the regulation of imprinted genes by showing that they contain PHE1 DNA-binding sites. Our data favour a scenario where these elements have been domesticated to function as providers of *cis*-regulatory sequences that facilitate transcription. Similarly, binding sites for type II MADS-box TFs, as well as for E2F TFs, have been shown to be carried by TEs (Hénaff et al., 2014; Muiño et al., 2016). These results, together with our study, provide examples of TE-mediated distribution of TF binding sites throughout flowering plant genomes, adding support to the long-standing idea that transposition facilitates the formation of *cis*-regulatory architectures required to control complex biological processes (Britten and Davidson, 1971; Feschotte, 2008; Hirsch and Springer, 2017). We speculate that this process may have contributed to endosperm evolution by allowing the recruitment of crucial developmental genes into a single transcriptional network, regulated by type I MADS-box TFs. The diversification of the mammalian placenta has been connected with the dispersal of hundreds of placenta-specific enhancers by endogenous retroviruses (Chuong et al., 2013; Dunn-Fletcher et al., 2018), suggesting that the convergent evolution of the endosperm in flowering plants and the mammalian placenta have been promoted by TE transpositions.

**Figure 1 – Figure Supplement 1.**
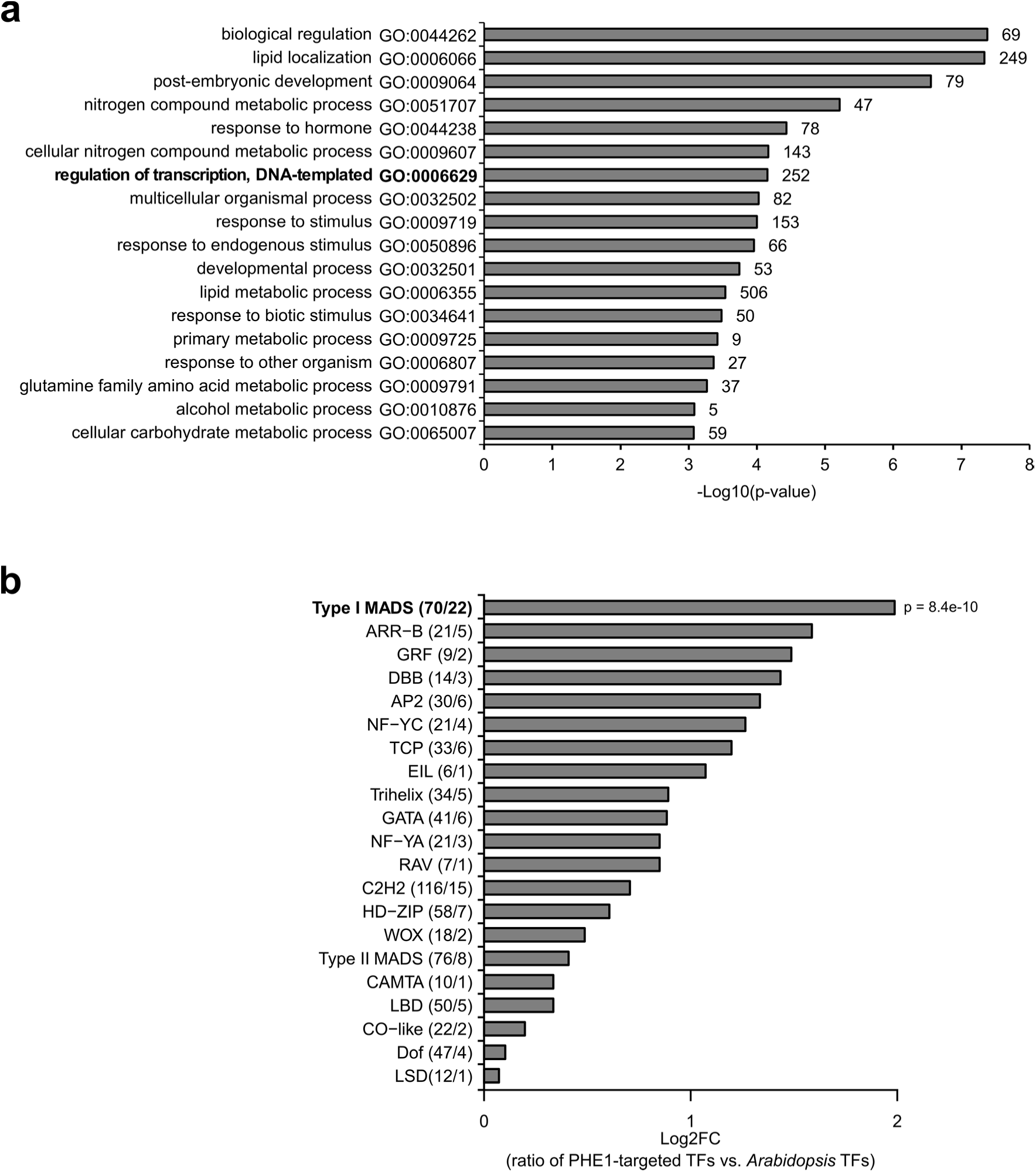
Transcription factor genes are enriched among PHE1 targets. **a,** Enriched biological processes within PHE1 target genes. Numbers on bars indicate number of PHE1 target genes within each GO-term. **b,** Enrichment of transcription factor (TF) families among PHE1 target genes (see Methods). Numbers in parenthesis indicate the total number of *Arabidopsis* genes belonging to a certain transcription factor family, and the total number of genes in that family targeted by PHE1, respectively. P-value was determined using the hypergeometric test.

**Figure 1 – Figure Supplement 2.**
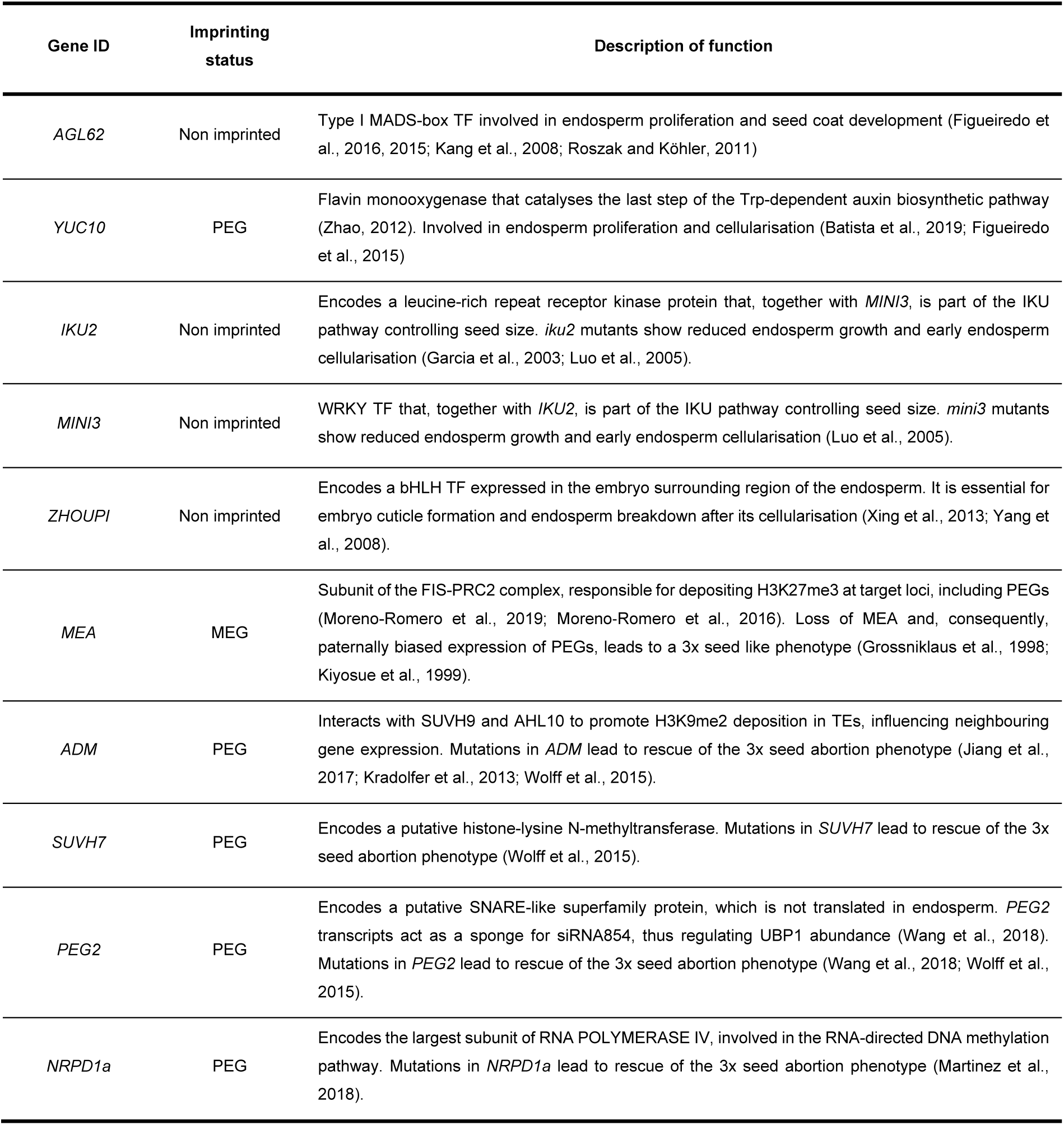
PHE1 target genes previously implicated in endosperm development.

**Figure 1 – Figure Supplement 3.**
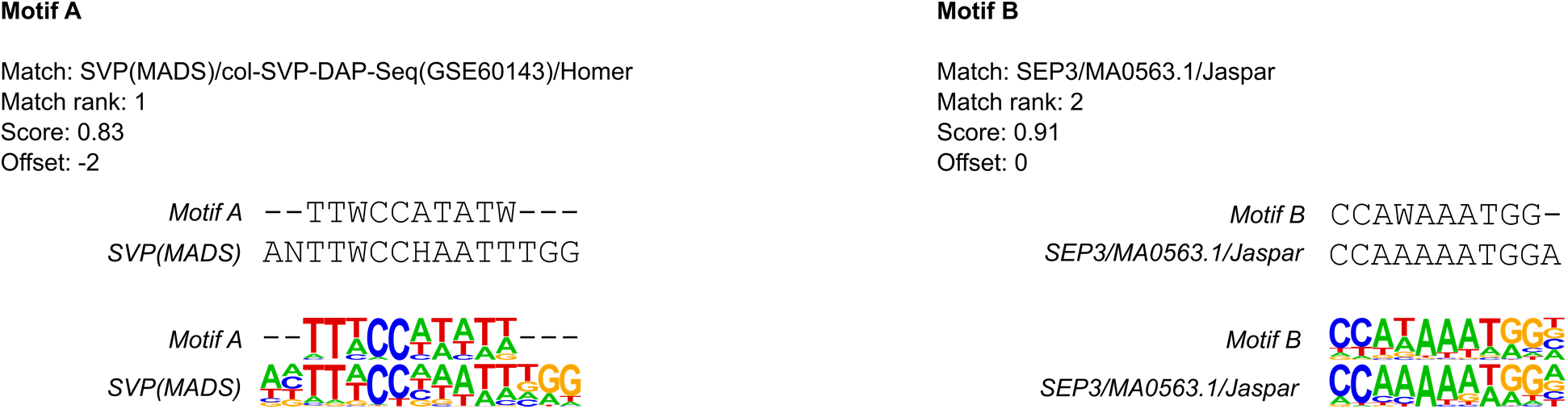
PHE1 DNA-binding motifs show similarity to type II MADS-box CArG-boxes. Alignment between PHE1 DNA-binding motifs and known motif matches.

**Figure 1 – Figure Supplement 4.**
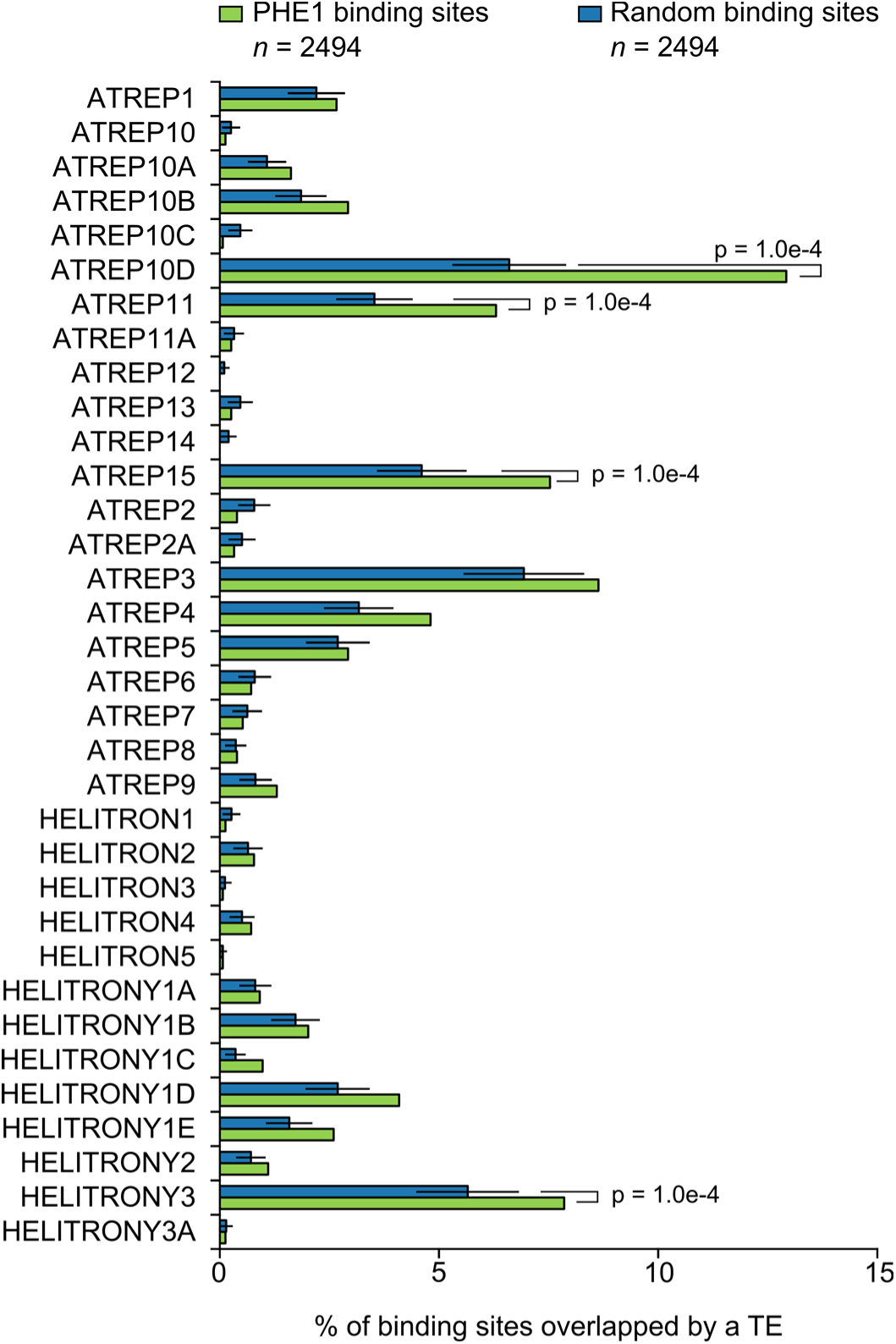
RC/Helitron family overlap with PHE1 binding sites. Fraction of PHE1 binding sites (green) overlapping with different RC/Helitron TE families. Overlap is expressed as the percentage of binding sites where spatial intersection with the families specified on the y-axis is observed. A set of random binding sites is used as control (blue). This control set was obtained by randomly shuffling the PHE1 binding sites in *A. thaliana* gene promoters (see Methods). P-values were determined using Monte Carlo permutation tests (see Methods). Bars represent ± sd, (*n* = 2494, PHE1 binding sites, Random binding sites).

**Figure 1 – Figure Supplement 5.**
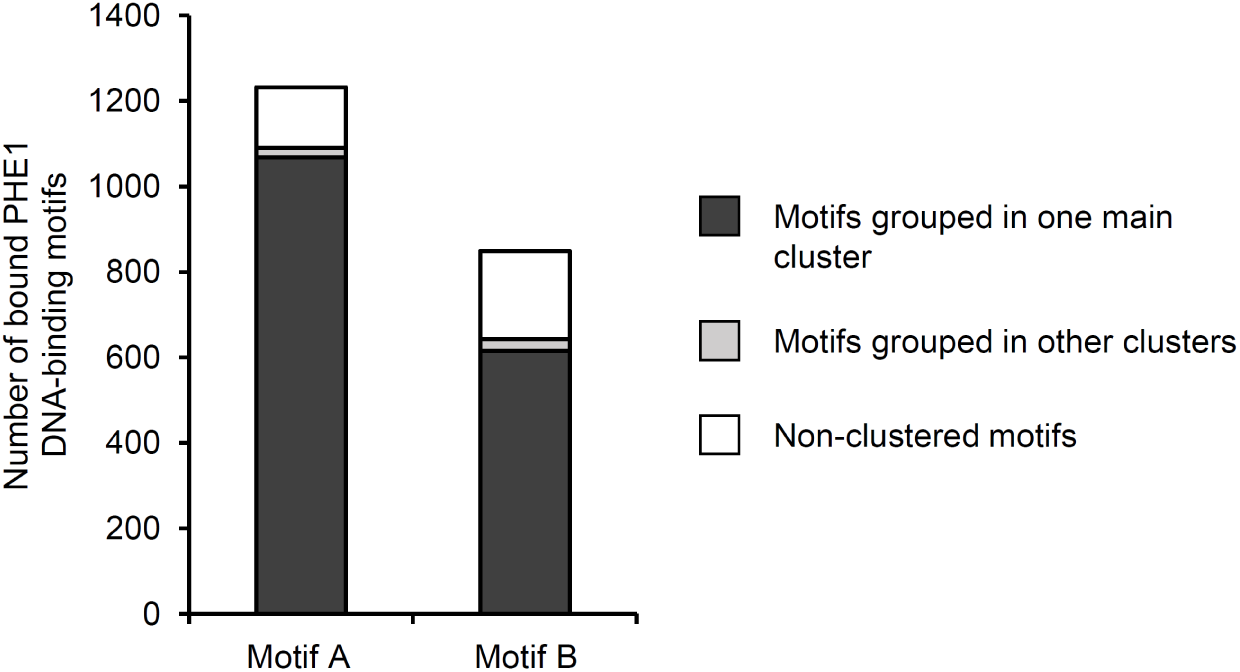
Clustering of bound PHE1 DNA-binding motifs. PHE1-DNA binding motifs and their flanking sequences were paired based on sequence homology. The pairwise homologous sequences were then merged into higher order clusters, based on shared elements in the homologous pairs (see Methods). Motifs are those identified in Fig. 1a.

**Figure 1 – Figure Supplement 6.**
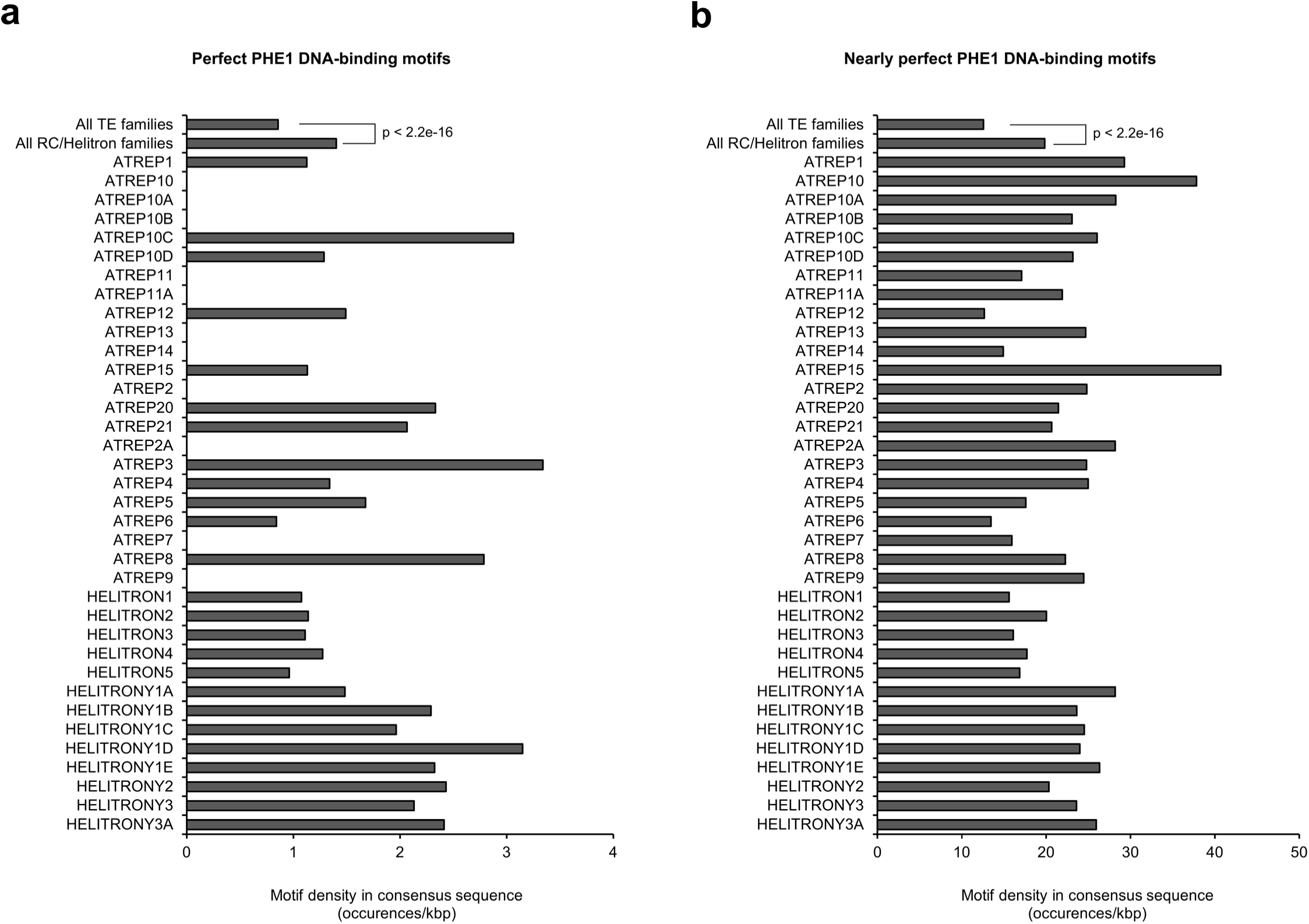
PHE1 DNA-binding motif densities in RC/Helitron consensus sequences. **a,** Density of perfect PHE1-DNA binding motifs in consensus sequences of different RC/Helitron families. Perfect motifs are those described in Fig. 1a. As a reference, the density of perfect PHE1 DNA-binding motifs in consensus sequences of all TE families is shown. **b,** Density of nearly perfect PHE1-DNA binding motifs in the consensus sequences of different RC/Helitron families. Nearly perfect motifs are those sequences where only one nucleotide substitution is required to generate a perfect PHE1 DNA-binding motif. As a reference, the density of nearly perfect PHE1 DNA-binding motifs in the consensus sequences of all TE families is shown. P-values were determined using χ^2^ tests.

**Figure 2 – Figure Supplement 1.**
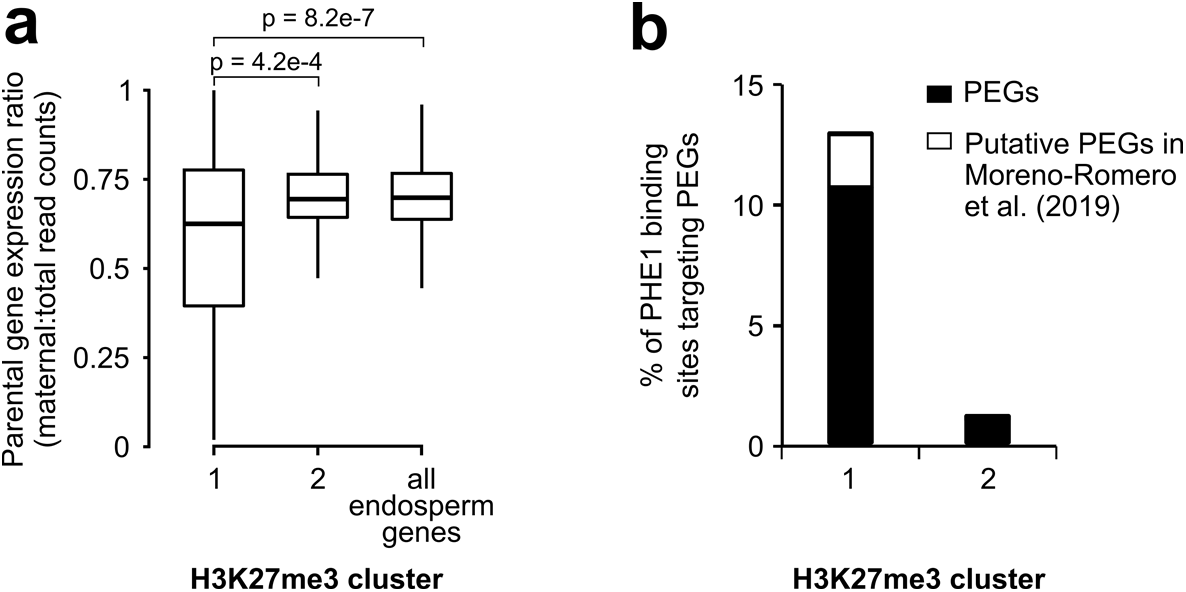
Characterization of H3K27me3 clusters. **a,** Parental gene expression ratio of PHE1 targets associated with Cluster 1 and Cluster 2 binding sites, and all endosperm expressed genes. P-values were determined using two-tailed Mann-Whitney tests. **b,** Fraction of Cluster 1 and Cluster 2 binding sites that target PEGs (**Fig. 1 – source data 1**) and putative PEGs (Moreno-Romero et al., 2019). (a-b) H3K27me3 clusters are those defined in Fig. 2b.

**Figure 2 – Figure Supplement 2.**
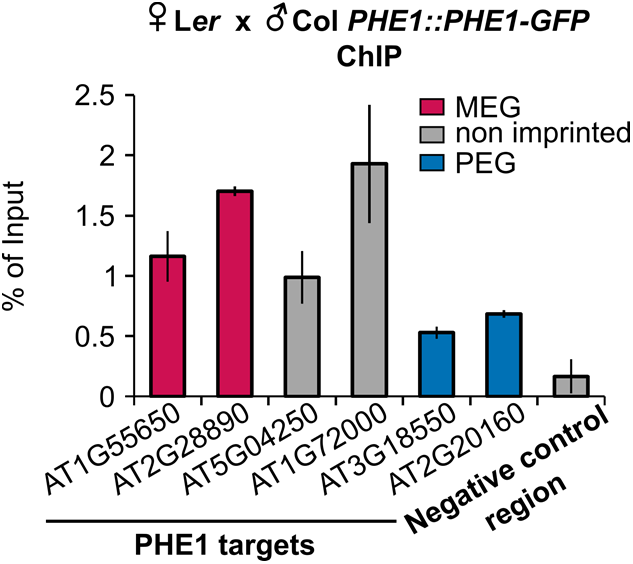
Parental-specific PHE1 ChIP. qPCR of purified ChIP-DNA. Enrichment is shown as % of input DNA, in regions associated with MEGs (pink), PEGs (blue), and non-imprinted (grey) PHE1 target genes. Bars represent ± sd. Data from one representative biological replicate is shown (*n =* 2 biological replicates).

**Figure 2 – Figure Supplement 3.**
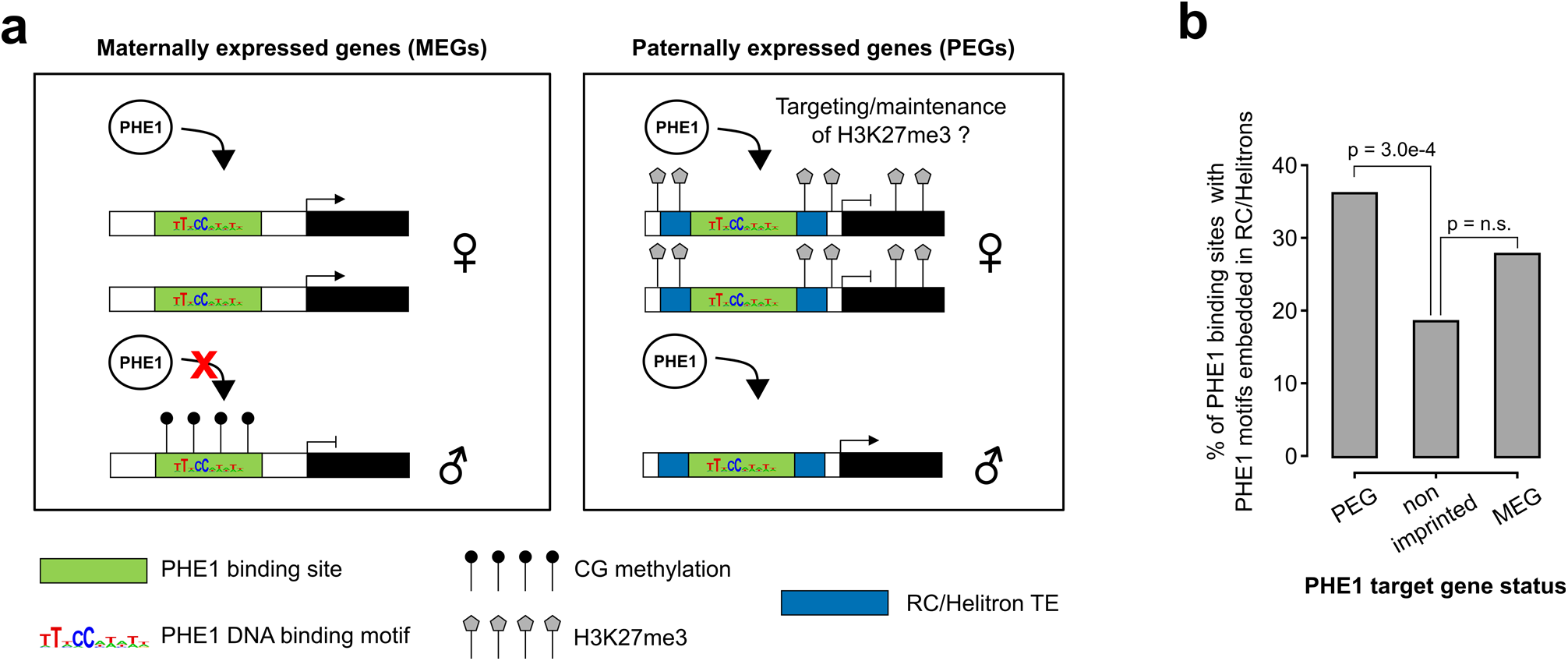
Control of imprinted gene expression by PHE1. **a,** Schematic model of imprinted gene control by PHE1. Maternally expressed genes (left panel) show DNA hypermethylation of paternal (♂) PHE1 binding sites. This precludes PHE1 accessibility to paternal alleles, leading to predominant binding and transcription from maternal (♀) alleles. In paternally expressed genes (right panel), RC/Helitrons found in flanking regions carry PHE1 DNA-binding motifs, allowing PHE1 binding. The paternal PHE1 binding site is devoid of repressive H3K27me3, facilitating binding of PHE1 and transcription of this allele. H3K27me3 accumulates at the flanks of maternal PHE1 binding sites, while the binding sites remain devoid of this repressive mark. PHE1 is able to bind maternal alleles, but fails to induce transcription. We hypothesize that the accessibility to maternal PHE1 binding sites might be important for deposition of H3K27me3 during central cell development (possibly by another type I MADS-box transcription factor). It may furthermore be required for maintenance of H3K27me3 during endosperm division. **b,** Fraction of PHE1 binding sites targeting MEGs, PEGs, or non-imprinted genes where a spatial overlap between a RC/Helitron and a PHE1 DNA-binding motif is observed (as illustrated in a, right panel). P-values were determined using the hypergeometric test.

**Figure 3 – Figure Supplement 1.**
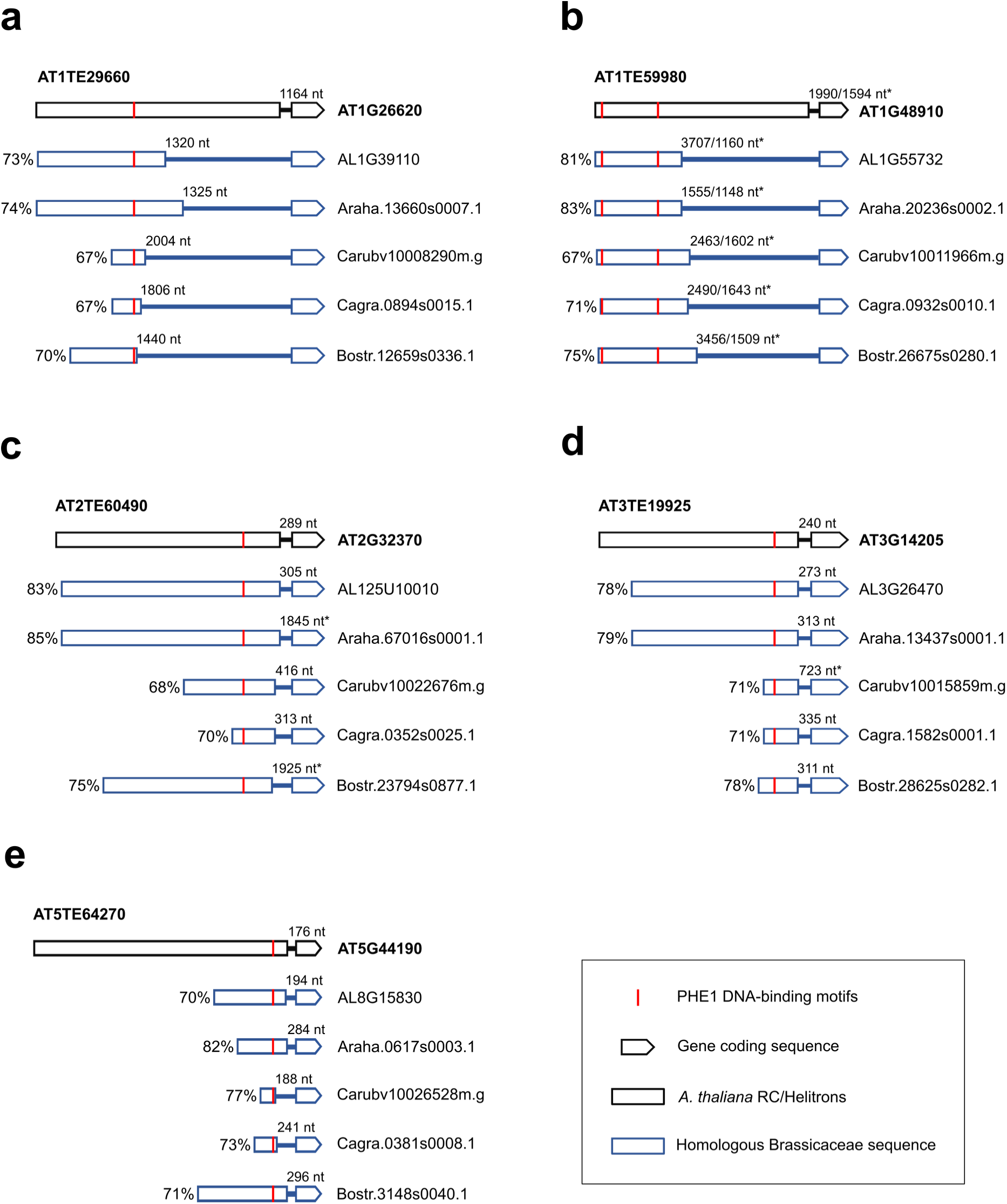
Homology analysis of PEGs and upstream RC/Helitron sequences within Brassicaceae. Alignments between *Arabidopsis thaliana* RC/Helitrons associated with PEGs (Fig. 3a-e) and homologous sequences in other Brassicaceae, where the putative homologous PHE1-DNA binding motifs are present across all six species. For each aligned fragment, the percent sequence identity is indicated. The distances between the DNA-binding motifs and the TSS of PEGs are labelled. * corresponds to the distance to the translation start site, in cases where the annotation for the 5’-UTR of the gene is not available.

**Figure 4 – Figure Supplement 1.**
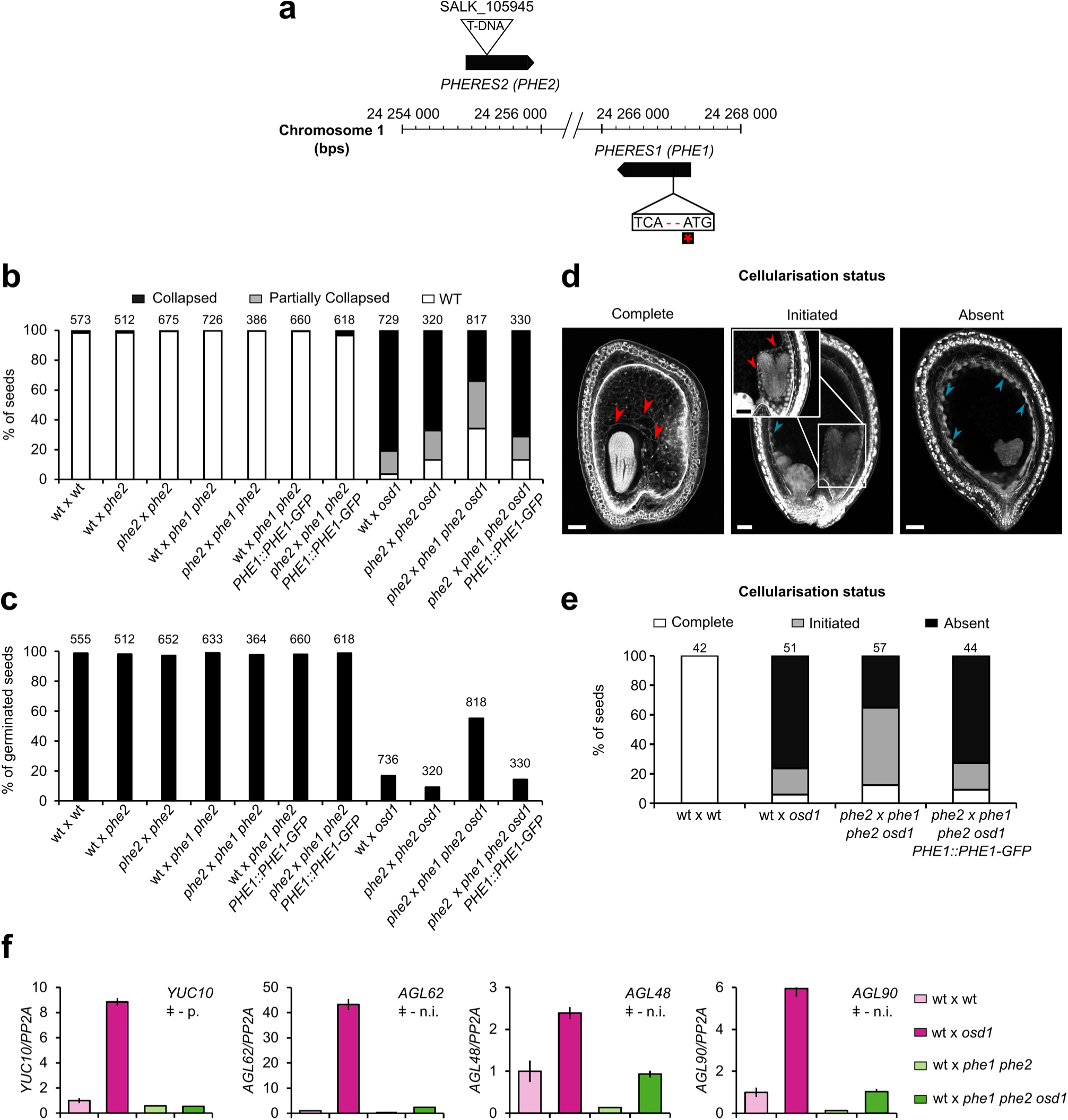
Rescue of 3x seed inviability in *phe1 phe2*. **a,** Schematic representation of the *phe1 phe2* mutant. CRISPR/Cas9 was used to generate a premature stop codon in *PHE1*, represented by the red asterisk. *PHE1* mutagenesis was done in the *phe2* background, due to the proximity and likely redundancy of both genes (see Methods). **b-c,** Seed inviability phenotype (b) of wild-type (wt) and paternal excess crosses, in wt, *phe1 phe2*, and *phe1* complementation lines, with respective seed germination rates (c). **d-e**, Status of endosperm cellularisation in wt and paternal excess seeds. (b-c,e) *n =* numbers on top of bars (seeds). **f,** Expression of PHE1 target genes in rescued 3x seeds. ǂ - gene neighbours an RC/Helitron containing a PHE1 DNA-binding motif; p. – PEG; n.i. – non-imprinted gene. Bars represent ± sd. Data from one representative biological replicate is shown (*n =* 2 biological replicates).

**Figure 4 – Figure Supplement 2.**
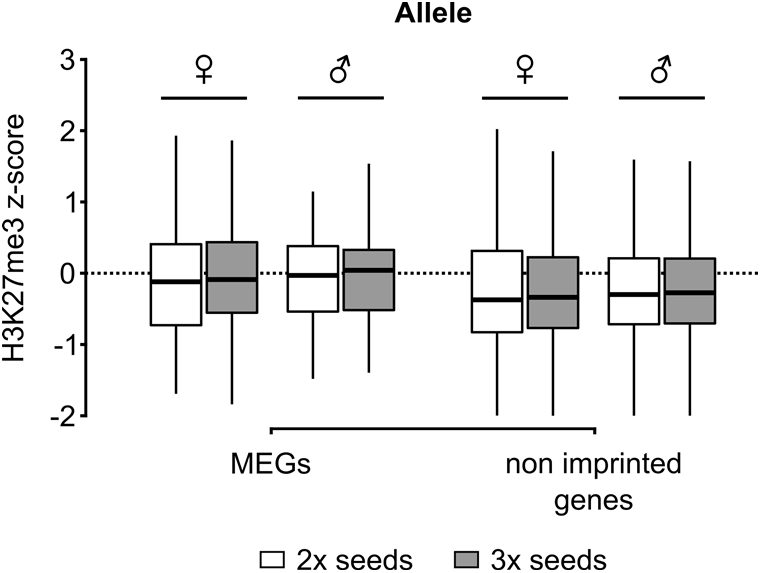
Distribution of H3K27me3 in 2x and 3x seeds. Accumulation of H3K27me3 in maternal (♀) and paternal (♂) gene bodies of MEGs and non-imprinted genes in the endosperm of 2x and 3x seeds (white and grey, respectively).

## Methods

### Plant material and growth conditions

*Arabidopsis thaliana* seeds were sterilized in a closed vessel containing chlorine gas, for 3 hours. Chlorine gas was produced by mixing 3 mL HCl 37% and 100 mL of 100% commercial bleach. Sterile seeds were plated in ½ MS-medium (0.43% MS salts, 0.8% Bacto Agar, 0.19% MES hydrate) supplemented with 1% Sucrose. When required, appropriate antibiotics were supplemented to the medium. Seeds were stratified for 48 h, at 4°C, in darkness. Plates containing stratified seeds were transferred to a long-day growth chamber (16h light / 8h dark; 110 μmol s^-1^m^-2^; 21°C; 70% humidity), where seedlings grew for 10 days. After this period, the seedlings were transferred to soil and placed in a long-day growth chamber.

Several mutant lines used in this study have been previously described: *osd1-1* (d’Erfurth et al., 2009), *osd1-3* (Heyman et al., 2011) and *pi-1* (Goto and Meyerowitz, 1994). The phe2 allele corresponds to a T-DNA insertion mutant (SALK_105945). Phenotypical analysis of this mutant revealed no deviant phenotype relative to Col wt plants (data not shown). Genotyping of phe2 was done using the following primers (PHE2 fw 5’-AAATGTCTGGTTTTATGCCCC-3’, PHE2 rv 5’-GTAGCGAGACAATCGATTTCG-3’, T-DNA 5’-ATTTTGCCGATTTCGGAAC-3’).

### Generation of phe1 phe2

The *phe1 phe2* double mutant was generated using the CRISPR/Cas9 technique. A 20-nt sgRNA targeting *PHE1* was designed using the CRISPR Design Tool (Ran et al., 2013). A single-stranded DNA oligonucleotide corresponding to the sequence of the sgRNA, as well as its complementary oligonucleotide, were synthesized. BsaI restriction sites were added at the 5’ and 3’ ends, as represented by the underlined sequences (sgRNA fw 5’-<u>ATTG</u>CTCCTGGATCGAGTTGTAC-3’; sgRNA rv 5’-<u>AAAC</u>GTACAACTCGATCCAGGAG-3’). These two oligonucleotides were then annealed to produce a double stranded DNA molecule.

The double-stranded oligonucleotide was ligated into the egg-cell specific pHEE401E CRISPR/Cas9 vector (Z.-P. Wang et al., 2015) through the BsaI restriction sites. This vector was transformed into the *Agrobacterium tumefaciens* strain GV3101, and *phe2/-* plants were subsequently transformed using the floral-dip method (Clough and Bent, 1998).

To screen for T1 mutant plants, we performed Sanger sequencing of *PHE1* amplicons derived from these plants, and obtained with the following primers (fw 5’-AGTGAGGAAAACAACATTCACCA-3’; rv 5’-GCATCCACAACAGTAGGAGC-3’). The selected mutant contained a homozygous two base pair deletion that leads to a premature stop codon, and therefore a truncated PHE1_1-50aa_ protein. In the T2 generation, the segregation of pHEE401E allowed to select plants that did not contain this vector and that were double homozygous *phe1 phe2* mutants. Genotyping of the *phe1* allele was done using primers fw 5’-AAGGAAGAAAGGGATGCTGA-3’ and rv 5’-TCTGTTTCTTTGGCGATCCT-3’, followed by RsaI digestion.

### Seed imaging

Analysis of endosperm cellularisation status was done following the Feulgen staining protocol described previously (Batista et al., 2019). Imaging of Feulgen-stained seeds was done in a Zeiss LSM780 NLO multiphoton microscope with excitation wavelength of 800 nm, and acquisition between 520 nm – 695 nm.

### RT-qPCR

RNA extraction, cDNA synthesis and qPCR analyses for *AGL* and *PEG* expression were performed as described previously (Batista et al., 2019). Two biological replicates per cross were used. Primer sequences for *YUC10*, *AGL62*, and *PP2A* were described previously (Batista et al., 2019). For the remaining genes, primer sequences were as follows: *AGL48* (fw 5’-TTCGCGATCCACCAGTGTTT-3’, rv 5’-GACCGCCTCCTACAAAACCA-3’), *AGL90* (fw 5’-TTGGTGATGAGTCGTTTTCCGA-3’, rv 5’-TCATATTCGCATTTGCGTCCG-3’).

### Chromatin immunoprecipitation (ChIP)

To find targets of PHE1, we performed a ChIP experiment using a reporter line containing the PHE1 protein tagged with GFP. This reporter line, which we denote as *PHE1::PHE1-GFP,* contains the *PHE1* promoter, its coding sequence and its 3’ regulatory sequence, and is present in a Col background (Weinhofer et al., 2010). Given the presence of the 3’ regulatory regions (Makarevich et al., 2008), this reporter behaves as a paternally expressed gene, similarly to the endogenous *PHE1* gene. Crosslinking of plant material was done by collecting 600 mg of 2 days after pollination (DAP) *PHE1::PHE1-GFP* siliques, and vacuum infiltrating them with a 1% formaldehyde solution in PBS. The vacuum infiltration was done for two periods of 15 min, with a vacuum release between each period. The crosslinking was then stopped by adding 0.125 mM glycine in PBS and performing a vacuum infiltration for a total of 15 min, with a vacuum release each 5 min. The material was then ground in liquid nitrogen, resuspended in 5mL Honda buffer (Moreno-Romero et al., 2017), and incubated for 15 min with gentle rotation. This mixture was filtered twice through Miracloth and one time through a CellTrics filter (30 µm), after which a centrifugation for 5 min, at 4°C and 1,500 g was performed. The nuclei pellet was then resuspended in 100 µL of nuclei lysis buffer (Moreno-Romero et al., 2017), and the ChIP protocol was continued as described before (Moreno-Romero et al., 2017). ChIP DNA was isolated using the Pure Kit v2 (Diagenode), following the manufacturer’s instructions.

For the parental-specific PHE1 ChIP the starting material consisted of 600 mg of 2DAP siliques from crosses between a *pi-1* mother (L*er* ecotype) and a *PHE1::PHE1-GFP* father (Col ecotype). The male sterile *pi-1* mutant was used to avoid emasculation of maternal plants. Crosslinking of plant material, nuclei isolation, ChIP protocol, and ChIP-DNA purification were the same as described before.

To assess parental-specific H3K27me3 profiles in 3x seeds, the INTACT system was used to isolate 4DAP endosperm of seeds derived from L*er pi-1* x Col *INTACT osd1-1* crosses, as described previously (Jiang et al., 2017; Moreno-Romero et al., 2017). *osd1* mutants were used for their ability to generate unreduced male gametes, leading to the formation of 3x seeds. ChIPs against H3 and H3K27me3 were then performed on the isolated endosperm nuclei, following the previously described protocol (Moreno-Romero et al., 2017).

The antibodies used for these ChIP experiments were as follows: GFP Tag Antibody (A-11120, Thermo Fisher Scientific), anti-H3 (Sigma, H9289), and anti-H3K27me3 (Millipore, cat. no. 07-449). All experiments were performed with two biological replicates.

### Library preparation and sequencing

*PHE1::PHE1-GFP* ChIP libraries were prepared using the Ovation Ultralow v2 Library System (NuGEN), with a starting material of 1 ng, following the manufacturer’s instructions. These libraries were sequenced at the SciLife Laboratory (Uppsala, Sweden), on an Illumina HiSeq2500 platform, using 50 bp single-end reads.

Library preparation and sequencing of H3K27me3 ChIPs in 3x seeds was done as described previously (Jiang et al., 2017).

Both datasets were deposited at NCBI’s Gene Expression Omnibus database (https://www.ncbi.nlm.nih.gov/geo/), under the accession number GSE129744.

### qPCR and Sanger sequencing of parental-specific PHE1 ChIP

Purified ChIP DNA and its respective input DNA obtained from the parental-specific PHE1 ChIP were used to perform qPCR. Positive and negative genomic regions for PHE1 binding were amplified using the following primers: AT1G55650 (fw 5’-CGAAGCGAAAAAGCACTCAC-3’; rv 5’-CCTTTTACATAATCCGCGTTAAA-3’), AT2G28890 (fw 5’-TTTGTGGTTGGAGGTTGTGA-3’; rv 5’-GTTGTTCGTGCCCATTTCTT-3’), AT5G04250 (fw 5’-AATTGACAAATGGTGTAATGGT-3’; rv 5’-CCAAAGAATTTGTTTTTCTATTCC-3’), AT1G72000 (fw 5’-AACAAATATGCACAAGAAGTGC-3’; rv 5’-ACCTAGCAAGCTGGCAAAAC-3’), AT3G18550 (fw 5’-TCCTTTTCCAAATAAAGGCATAA-3’; rv 5’-AAATGAAAGAAATAAAAGGTAATGAGA-3’), AT2G20160 (fw 5’-TCCTAAATAAGGGAAGAGAAAGCA-3’; rv 5’-TGTTAGGTGAAACTGAATCCAA-3’), negative region (fw 5’-TGGTTTTGCTGGTGATGATG-3’, rv 5’-CCATGACACCAGTGTGCCTA-3’). HOT FIREPol EvaGreen qPCR Mix Plus (ROX) (Solis Biodyne) was used as a master mix for qPCR amplification in a iQ5 qPCR system (Bio-Rad).

For Sanger sequencing, positive genomic regions for PHE1 binding containing SNPs that allowed distinction between parents were amplified by PCR using the Phusion High-Fidelity DNA Polymerase (Thermo Fisher Scientific), in combination with the primers described above. Amplified DNA was purified using the GeneJET PCR Purification kit (Thermo Fisher Scientific) and used for Sanger sequencing. The chromatograms obtained from Sanger sequencing were then analysed for the presence of SNPs. The maternal:total read ratios were retrieved from the publications where these genes were identified as imprinted.

### Bioinformatic analysis of ChIP-seq data

For the *PHE1::PHE1-GFP* ChIP, reads were aligned to the *Arabidopsis* (TAIR10) genome using Bowtie version 1.2.2 (Langmead, 2010), allowing 2 mismatches (-v 2). Only uniquely mapped reads were kept. ChIP-seq peaks were called using MACS2 version 2.1.1, with its default settings (--gsize 1.119e8, --bw 250) (Zhang et al., 2008). Input samples served as control for their corresponding GFP ChIP sample. Each biological replicate was handled individually using the same peak calling settings, and only the overlapping peak regions between the two replicates were considered for further analysis. These regions are referred throughout the text as PHE1 binding sites (**Supplementary File 1**). Peak overlap was determined with BEDtools version 2.26.0 (Quinlan and Hall, 2010). Each PHE1 binding site was annotated to a genomic feature and matched with a target gene using the peak annotation feature (annotatePeaks.pl) provided in HOMER version 4.9 (Heinz et al., 2010) (**Supplementary File 1**). Only binding sites located less than 3 kb away from the nearest transcription start site were considered.

PHE1 DNA-binding motifs were identified from *PHE1::PHE1-GFP* ChIP-seq peak regions with HOMER’s findMotifsGenome.pl function, using the default settings. P-values of motif enrichment, as well as alignments between PHE1 motifs and known motifs were generated by HOMER.

Read mapping, coverage analysis, purity calculations, normalisation of data, and determination of parental origin of reads derived from H3K27me3 ChIPs in 3x seeds was done following previously published methods (Moreno-Romero et al., 2016) (**Supplementary File 1**).

### Analysis of PHE1 target genes

Significantly enriched Gene Ontology terms within target genes of PHE1 were identified using AtCOECIS (Vandepoele et al., 2009), and further summarized using REVIGO (Supek et al., 2011).

Enrichment of specific transcription factor families within PHE1 targets was calculated by first normalizing the number of PHE1-targeted TFs in each family, to the total number of TFs targeted by PHE1. As a control, the number of TFs belonging to a certain family was normalised to the total number of TFs in the *Arabidopsis* genome. The Log_2_ fold change between these ratios was then calculated for each family. Significance of the enrichment was assessed using the hypergeometric test. Annotation of transcription factor families was done following the Plant Transcription Factor Database version 4.0 (Jin et al., 2014). Only TF families containing more than 5 members were considered in this analysis.

To determine which imprinted genes are targeted by PHE1, a custom list consisting of the sum of imprinted genes identified in different studies was used (**Fig. 1 – source data 1**) (Gehring et al., 2011; Hsieh et al., 2011; Pignatta et al., 2014; Schon and Nodine, 2017; Wolff et al., 2011).

To determine the proportion of genes overexpressed in paternal excess crosses that are targeted by PHE1, a previously published transcriptome dataset of 3x seeds was used (Schatlowski et al., 2014).

### Spatial overlap of TEs and PHE1 binding sites

Spatial overlap between PHE1 ChIP-seq peak regions (binding sites) and TEs was determined using the regioneR package version 1.8.1 (Gel et al., 2015), implemented in R version 3.4.1 (Core Team R, 2017). As a control, a mock set of binding sites was created, to which we refer to as random binding sites. This random binding site set had the same total number of binding sites and the same size distribution as the PHE1 binding site set. Using regioneR, a Monte Carlo permutation test with 10000 iterations was performed. In each iteration the random binding sites were arbitrarily shuffled in the 3 kb promoter region of all *Arabidopsis thaliana* genes. From this shuffling, the average overlap and standard deviation of the random binding site set was determined, as well as the statistical significance of the association between PHE1 binding sites and TE superfamilies/families.

BedTools version 2.26.0 (Quinlan and Hall, 2010) was used to determine the fraction of PHE1 binding sites targeting MEGs, PEGs, or non-imprinted genes where a spatial overlap between binding sites, RC/Helitrons and PHE1 DNA-binding motifs is simultaneously observed. The hypergeometric test was used to assess the significance of the enrichment of PHE1 binding sites where this overlap is observed, across different target types.

### Expression analyses of genes flanked by RC/Helitrons

We investigated the expression level of genes containing RC/Helitrons in their promoter regions (defined as 3kb upstream of the TSS), in embryo, endosperm, and seed coat of pre-globular stage seeds. Affymetrix GeneChip ATH1 Arabidopsis Genome Array data were extracted from Belmonte et al., (2013). The expression values in micropylar endosperm, peripheral endosperm, and chalazal endosperm were averaged to represent the endosperm expression level. The endosperm expression levels of a given gene were then compared to the expression levels in the embryo and seed coat.

RC/Helitron-associated genes were classified into three groups: (i) genes with RC/Helitrons and PHE1 DNA-binding motifs within the 3kb promoter, thus representing genes with domesticated RC/Helitrons; (ii) genes having RC/Helitrons and PHE1 DNA-binding motifs within the 3kb promoter, but not bound by PHE1; and (iii) genes having RC/Helitrons located within the 3kb promoter, and no PHE1 DNA-binding motifs. A two-tailed Mann-Whitney test with continuity correction was used to assess statistical significance of differences in expression levels between gene groups.

### Calculation of PHE1 DNA-binding motif densities

To measure the density of PHE1 DNA-binding motifs within different genomic regions of interest, the fasta sequences of these regions were first obtained using BEDtools. HOMER’s scanMotifGenomeWide.pl function was then used to screen these sequences for the presence of PHE1 DNA-binding motifs, and to count the number of occurrences of each motif. Motif density was then calculated as the number of occurrences of each motif, normalized to the size of the genomic region of interest. Motif densities in RC/Helitron consensus sequences were calculated as above. Perfect PHE1 DNA-binding motif sequences were defined as those represented in Fig. 1a. Nearly perfect motif sequences were defined as sequences where only one nucleotide substitution could give rise to a perfect PHE1 DNA-binding motif. Consensus sequences were obtained from Repbase (Bao et al., 2015). Chi-square tests of independence were used to test if there were any associations between specific genomic regions and PHE1 DNA-binding motifs. This was done by comparing the proportion of DNA bases corresponding to PHE1 DNA-binding motifs in each genomic region.

### Identification of homologous PHE1 DNA-binding motifs carried by RC/Helitrons

To assess the homology of PHE1 DNA-binding motifs and associated RC/Helitron sequences, pairwise comparisons were made among all sequences, using the BLASTN program. The following parameters were followed: word size = 7, match/mismatch scores = 2/-3, gap penalties, existence = 5, extension = 2. The RC/Helitron sequences were considered to be homologous if the alignment covered at least 9 out of the 10 bp PHE1 DNA-binding motif sites, extended longer than 30 bp, and had more than 70% identity. Because the mean length of intragenomic conserved non-coding sequences is around 30 bp in *A. thaliana* (Thomas et al., 2007), we considered this as the minimal length of alignments to define a pair of related motif-carrying TE sequences. The pairwise homologous sequences were then merged in higher order clusters, based on shared elements in the homologous pairs.

### Phylogenetic analyses of PHE1-targeted PEG orthologs in the Brassicaceae

Amino acid sequences and nucleotide sequences of PHE1-targeted PEGs were obtained from TAIR10. The sequences of homologous genes in the Brassicaceae and several other rosids were obtained in PLAZA 4.0 (https://bioinformatics.psb.ugent.be/plaza/) (Van Bel et al., 2018), BRAD database (http://brassicadb.org/brad/) (X. Wang et al., 2015), and Phytozome v.12 (https://phytozome.jgi.doe.gov/) (Goodstein et al., 2012).

For each PEG of interest, the amino acid sequences of the gene family were used to generate a guided codon alignment by MUSCLE with default settings (Edgar, 2004). A maximum likelihood tree was then generated by IQ-TREE 1.6.7 with codon alignment as the input (Nguyen et al., 2015). The implemented ModelFinder was executed to determine the best substitution model (Kalyaanamoorthy et al., 2017), and 1000 replicates of ultrafast bootstrap were applied to evaluate the branch support (Hoang et al., 2018). The tree topology and branch supports were reciprocally compared with, and supported by another maximum likelihood tree generated using RAxML v. 8.1.2 (Stamatakis, 2014).

We selected PEGs that had well supported gene family phylogeny with no lineage-specific duplication in the *Arabidopsis* and *Capsella* clades, and where imprinting data were available for all *Capsella grandiflora* (Cgr), *C. rubella* (Cru), and *Arabidopsis lyrata* (Aly) orthologs of interest (Klosinska et al., 2016; Lafon-Placette et al., 2018; Pignatta et al., 2014). We then obtained the promoter region, defined as 3 kb upstream of the TSS, of the orthologs and paralogs in Brassicaceae and rosids species. These promoter sequences were searched for the presence of homologous RC/Helitron sequences, as well as for putative PHE1 DNA-binding sites contained in these TEs.

Homology between *A. thaliana* RC/Helitron sequences and Brassicaceae sequences was detected by aligning the *A. thaliana* sequence to the promoter region of the orthologous PEG, using the BLASTN program, with the following parameters: word size = 11, match/mismatch scores = 2/-3, gap penalties, existence = 5, extension = 2. Aligned sequences are considered homologous if spanning more than 100 nt of the *Arabidopsis thaliana* RC/Helitron, with over 60% identity.

### Epigenetic profiling of PHE1 binding sites

Parental-specific H3K27me3 profiles (Moreno-Romero et al., 2016) and DNA methylation profiles (Schatlowski et al., 2014) generated from endosperm of 2x seeds were used for this analysis. Levels of H3K27me3 and CG DNA methylation were quantified in each 50 bp bin across the 2 kb region surrounding PHE1 binding site centres using deepTools version 2.0 (Ramírez et al., 2016). These values were then used to generate H3K27me3 heatmaps and metagene plots, as well as boxplots of CG methylation in PHE1 binding sites. Clustering analysis of H3K27me3 distribution in PHE1 binding sites was done following the k-means algorithm as implemented by deepTools. A two-tailed Mann-Whitney test with continuity correction was used to assess statistical significance of differences in CG methylation levels.

### Parental gene expression ratios in 2x and 3x seeds

To determine parental gene expression ratios in 2x and 3x seeds, we used previously generated endosperm gene expression data (Martinez et al., 2018). In this dataset, L*er* plants were used as maternal plants pollinated with wt Col or *osd1* Col plants, allowing to determine the parental origin of sequenced reads following the method described before (Moreno-Romero et al., 2019).

Parental gene expression ratios were calculated as the number of maternally-derived reads divided by the sum of maternally- and paternally-derived reads available for any given gene. Ratios were calculated separately for the two biological replicates of each cross (L*er* x wt Col and L*er* x *osd1* Col), and the average of both replicates was considered for further analysis. The MEG and PEG ratio thresholds for 2x and 3x seeds indicated in Figure 3d were defined as a four-fold deviation of the expected read ratios, towards more maternal or paternal read accumulation, respectively. The expected read ratio for a biallelically expressed gene in 2x seeds is 2 maternal reads : 3 total reads, while for 3x paternal excess seeds this ratio is 2 maternal reads : 4 total reads. Deviations from these expected ratios were used to classify the expression of published imprinted genes (**Fig. 1 – source data 1**) as maternally or paternally biased in 3x seeds, according to the direction of the deviation. As a control, the parental bias of these imprinted genes was also assessed in 2x seeds.

### Parental expression ratios of genes associated with H3K27me3 clusters

Previously published endosperm gene expression data, generated with the INTACT system, was used for this analysis (Del Toro-De León and Köhler, 2018). Parental gene expression ratios were determined as the mean between ratios observed in the L*er* x Col cross and its reciprocal cross. As a reference, the parental gene expression ratio for all endosperm expressed genes was also determined. A two-tailed Mann-Whitney test with continuity correction was used to assess statistical significance of differences between parental gene expression ratios.

### H3K27me3 accumulation in imprinted genes

Parental-specific accumulation of H3K27me3 across imprinted gene bodies in the endosperm of 2x (Moreno-Romero et al., 2016) and 3x seeds (this study) was estimated by calculating the mean values of the H3K27me3 z-score across the gene length. Imprinted genes were considered as those genes previously identified in different studies (Gehring et al., 2011; Hsieh et al., 2011; Pignatta et al., 2014; Schon and Nodine, 2017; Wolff et al., 2011). A two-tailed Mann-Whitney test with continuity correction was used to assess statistical significance of differences in H3K27me3 z-score levels.

### Statistics

Sample size, statistical tests used, and respective p-values are indicated in each figure or figure legend, and further specified in the corresponding Methods sub-section.

### Data availability

ChIP-seq data generated in this study is available at NCBI’s Gene Expression Omnibus database (https://www.ncbi.nlm.nih.gov/geo/), under the accession number GSE129744. Additional data used to support the findings of this study are available at NCBI’s Gene Expression Omnibus, under the following accession numbers: H3K27me3 ChIP-seq data from 2x endosperm (Moreno-Romero et al., 2016) - GSE66585; Gene expression data in 2x and 3x endosperm (Martinez et al., 2018) - GSE84122; Gene expression data in 2x and 3x seeds and parental-specific DNA methylation from 2x endosperm (Schatlowski et al., 2014) - GSE53642; Parental-specific gene expression data of 2x INTACT-isolated endosperm nuclei (Del Toro-De León and Köhler, 2018) - GSE119915. Gene expression profile of different seed compartments (Belmonte et al., 2013) – GSE12404.

## Acknowledgments

We thank Qi-Jun Chen for providing the pHEE401E CRISPR/Cas9 vector. We are grateful to Cecilia Wärdig for technical assistance. Sequencing was performed by the SNP&SEQ Technology Platform, Science for Life Laboratory at Uppsala University, a national infrastructure supported by the Swedish Research Council (VRRFI) and the Knut and Alice Wallenberg Foundation. This research was supported by a grant from the Swedish Research Council (to C.K.), a grant from the Knut and Alice Wallenberg Foundation (to C.K.), and support from the Göran Gustafsson Foundation for Research in Natural Sciences and Medicine.

## Author contributions

R.A.B., J.M-R., Y.Q, D.D.F and C.K. performed the experimental design. R.A.B., J.M-R., J.V.B. and Y.Q. performed experiments. R.A.B., J.M-R., Y.Q., J.S-G., and C.K. analysed the data. R.A.B and C.K. wrote the manuscript. All authors read and commented on the manuscript.

## Competing interests

The authors declare no competing interests.

## Materials and correspondence

The materials generated in this study are available upon request to C.K.

